# Attention rhythmically shapes sensory tuning

**DOI:** 10.1101/2023.05.26.542055

**Authors:** Laurie Galas, Ian Donovan, Laura Dugué

## Abstract

Attention is key to perception and human behavior, and evidence shows that it periodically samples sensory information (<20Hz). However, this view has been recently challenged due to methodological concerns and gaps in our understanding of the function and mechanism of rhythmic attention. Here we used an intensive ∼22-hour psychophysical protocol combined with reverse correlation analyses to infer the neural representation underlying these rhythms. Participants performed a task in which covert spatial (sustained and exploratory) attention was manipulated, and then probed at various delays. Our results show that sustained and exploratory attention periodically modulate perception via different neural computations. While sustained attention suppresses distracting stimulus features at the alpha (∼12Hz) frequency, exploratory attention increases the gain around task-relevant stimulus feature at the theta (∼6Hz) frequency. These findings reveal that both modes of rhythmic attention differentially shape sensory tuning, expanding the current understanding of the rhythmic sampling theory of attention.

**Significance statement:** For the past decade, low-frequency rhythms were described in attentional performance. Here, we go beyond description and assess the underlying neural computations in the sensory system. We used an intensive psychophysical protocol combined with reverse correlation analysis to infer the system’s sensitivity to and selectivity for stimulus feature (orientation) across time, and for two attention modes, i.e., sustained and exploratory attention. Our results show that signal enhancement as well as suppression occurred rhythmically, depending on the attentional state, leading to enhanced and improved performance phases – sustained and exploratory attention modes differentially shape the sensory tuning to stimulus feature.

## Introduction

Attention selectively facilitates sensory information processing, efficiently allocating resources based on task demands (Lennie, 2003; Carrasco, 2011). When voluntarily deployed, attention’s influence on perception can be sustained at will, benefiting performance for long periods of time. Although this classic understanding of sustained attention is well established, recent evidence suggests that attention periodically samples information, across both time and spatial locations (VanRullen, 2016; Fiebelkorn and Kastner, 2019; Kienitz et al., 2022).

Periodicity in behavioral performance has been assessed with paradigms including a sensory stimulus presented to reset participants’ attention at a specified time and spatial location (button press, saccade, arm movement are also used (Sauseng et al., 2007)), followed by a probe stimulus assessing changes in the spatio-temporal dynamics of the attention focus at various delays after the attentional reset. The temporal probing needs to be dense enough and across a long enough window for appropriate frequency exploration (Kienitz et al., 2022). Behavioral sampling studies have shown rhythmic fluctuations in performance at low frequency (<20Hz) due to attention (also observed for, e.g., expectation and serial dependence (Ho et al., 2022; Keitel et al., 2022)), suggesting that sensory information processing alternates between periods of greater (when in the attention focus) and lesser (when outside) facilitation (Landau and Fries, 2012; Fiebelkorn et al., 2013; Dugué and VanRullen, 2014; Song et al., 2014; Dugué et al., 2015b; Landau et al., 2015; Ho et al., 2017; Senoussi et al., 2019; Michel et al., 2022). Specifically, when attention is cued to a target location (valid cue) –sustained at one location preceding the probe– performance in the probe task fluctuates in alpha (∼12Hz). When attention is cued to a distractor stimulus location (invalid) –attention must shift and explore an uncued location– performance in the probe task fluctuates in theta (∼6Hz) (Dugué and VanRullen, 2017; Kienitz et al., 2022).

The view that attention modulates performance rhythmically has, however, been recently challenged (Ruzzoli et al., 2019; Keitel et al., 2022; van der Werf et al., 2022) notably due to methodological considerations (low trial number (van der Werf et al., 2022), low sampling frequency resolution (Kienitz et al., 2022) and issues regarding analyses (Brookshire, 2022); but see (Fiebelkorn, 2022a, 2022b; Re et al., 2022)). Additionally, while distinct frequency profiles of sustained and exploratory attention are consistent with distinct mechanisms and functional roles, the specific neural computations underlying variations in behavioral performance are still poorly understood. Specifically, it is unclear whether periodic changes in behavioral performance due to either attention mode are a result of fluctuations in signal enhancement or in noise suppression, two distinct neural computations by which attention shapes performance and sensory representations (Dosher and Lu, 2000; Lu and Dosher, 2000).

We used an intensive protocol for measuring periodic fluctuations of attentional performance in individual participants, and assessed how sensory tuning fluctuates with either exploratory or sustained attention. Following a pre-cue manipulating voluntary attention, a first discrimination task ensures exploratory and sustained attention were successfully manipulated. At various delays after the first task, a second detection task (vertical Gabor within noise) interrogates the dynamics of attentional stimulus sampling (Dugué et al., 2015b, 2017; Senoussi et al., 2019; Michel et al., 2022). Critically, reverse correlation analysis (Ahumada, 2002; Eckstein et al., 2002; Paltoglou and Neri, 2012; Wyart et al., 2012; Li et al., 2016; Tu et al., 2023; Xue et al., 2024) assessed potential fluctuations in sensory representations during the probe period. By regressing the stimulus energy across trials with behavioral responses in the detection task, we extracted orientation tuning curves of participants’ decision weights and evaluated signal enhancement and noise suppression over time for each attention mode. The results replicate previous findings of distinct rhythmic fluctuations of attentional performance for different attention modes (alpha for sustained and theta exploratory), and crucially reveal distinct sensory tuning mechanisms for either attention mode.

## Materials and Methods

### Participants

Nineteen participants were recruited for this experiment (11 female, age M ± SD = 23.26 ± 3.80 years; range 18-31). Two participants were excluded from the analyses because they did not complete the full experiment, two because they were not able to perform the task and four because they did not exhibit a significant attentional effect (see Analyses and Statistics). All participants had normal or corrected-to-normal vision and reported no history of psychiatric or neurological disorders, gave their written informed consent and were compensated for their participation. All procedures were approved by the French ethics committee Ouest IV - Nantes (IRB #20.04.16.71057) and followed the Code of Ethics of the World Medical Association (Declaration of Helsinki).

### Apparatus and stimuli

Participants sat in a dark room at 57cm from a calibrated and linearized CRT monitor (800*600 pixels, width = 37.8cm, height = 28.4cm, refresh rate 120Hz). Their heads were positioned on a chin rest to maintain the distance between the monitor and the eyes. Stimuli were generated using PsychToolbox 3 (3.0.12) (Brainard, 1997; Pelli, 1997; Kleiner et al., 2007) in Matlab R2014b (MathWorks). Background was gray (127.5, 127.5, 127.5 RGB). A black central fixation annulus with inner eccentricity of 0.2° (degree of visual angle) and thickness of 0.16° was presented throughout the experiment. The pre-cue and response cue were white rectangles (255, 255, 255 RGB, 0.5° long, 0.16° large) adjacent to the fixation (0.65° eccentricity from the center) and pointing down at a 45° angle toward the bottom left or bottom right. Stimuli were two conventional Landolt-C optotypes (2° high, 2° wide, 5.5° from the center at 45° angle toward the left or the right), and randomly generated circular noise patches at the same locations (2° diameter) with or without embedded Gabors (3 cycles per degree; circular window).

### Eye-tracking

The dominant eye of each participant was monitored using an infrared video camera (EyeLink 1000 plus, SR Research, Ottawa, Canada). Participants were instructed to maintain fixation. Each trial started contingent on fixating, and they had to maintain fixation until after the probe presentation. If a fixation break occurred, i.e., if participants’ gaze deviated by >1.5° or if they blinked, the trial was stopped and removed from the analysis. Supernumerary trials were added at the end of each block to compensate for rejected trials (M ± SD = 2257.73 ± 1320.76 fixation breaks total on average across participants).

### Procedure

Participants performed a total of 22 sessions. First, in a 1h-training session they were familiarized with the trial sequence at slower delays. They then performed two staircase blocks (see below) in an additional 1h-session. They finally performed 20 1h-long sessions (2 sessions per day with a break of 30 min between session; the entire experiment took on average 2.5 months) of the main task (totaling a number of 10,080 trials per participant).

#### Main task

Each session was composed of 4 blocks of 126 trials each as well as supernumerary trials in case of fixation break (see **Eye-tracking** section). Central fixation remained on the screen during the entire block. After a 1000ms-stable fixation a pre-cue appeared for 60ms followed by a 400ms-inter-stimulus interval (ISI). The two Landolt-Cs were displayed for 60ms (one in each lower quadrant) along with a response cue indicating which of the two was the target. Participants were instructed to discriminate the position of the C gap (randomly presented on the left or the right) of the target Landolt-C (their response was collected at the end of the trial sequence). When the response cue corresponded to the pre-cued location (2/3 of the trials), the trial was valid; when the response cue corresponded to the un-precued location (1/3 of the trials), the trial was invalid. Then a second, variable ISI (randomly chosen among 14 different possibilities equally distributed between 40ms and 495ms) preceded the presentation of two new stimuli: two patches of noise or noise+Gabor for 33ms, at the same location as the previous Landolt-Cs, followed by a 60ms delay. Participants were instructed to report the presence versus absence of the Gabor inside each patch of noise. The presence or absence of a Gabor in either patch within a trial was independent and random (50%), such that one, the other, both, or neither having a Gabor embedded were equally likely. These patches presented at multiples delays after the previous Landolt-Cs were used to probe the state of attention across time. At the end of the trial, participants were first asked to report the position of the gap in the target Landolt-C (press C key for a gap on the left with the left hand, and press B key for a gap on the right with the right hand; maximum response time of 1500ms), and then to report the presence or the absence of the Gabor inside each patch (for left patch, press C key for Gabor present and D key for Gabor absent with left hand; for right patch, press B key for Gabor present and H key for Gabor absent with right hand; maximum response time 2000ms).

#### Staircases

Two independent staircases were implemented via a Best PEST procedure (Pentland, 1980) using the Palamedes toolbox (Prins and Kingdom, 2018) in Matlab, to adapt first the size of the Landolt-C gap in the discrimination task, and second the SNR (signal- (Gabor) to-noise ratio) for the detection task. In the first staircase (208 trials), only the discrimination task sequence was presented and the size of the Landolt-Cs gap (1.4 ± 0.4° on average across participants) was adjusted according to a running-fit estimation (Best PEST) of the alpha parameter of a Weibull distribution, corresponding to the gap-size threshold reaching between 70% and 80% performance accuracy (76.97 ± 2.39% across participants in the main task). Here the trial sequence consisted of a neutral pre-cue (both the left and right pre-cue) for 60ms, a 400ms-ISI and the Landolt-Cs together with the response cue for 60ms. Participants had 1.5s to report the location of the gap. In the second staircase (200 trials), the procedure was the same. Only the detection task sequence was presented and the ratio between noise and Gabor was adapted to reach around 50% accuracy (62.64 ± 10.68 % across participants in the main task). One trial consisted of neutral pre-cue for 60ms, a 400ms-ISI and the 2 probe patches (50% noise only and 50% noise+Gabor, at each location) with a response cue for 33ms. Participants had 2s to report the presence or the absence of the Gabor in each of the two patches.

### Analyses and Statistics

Except when otherwise specified, all analyses were performed using Matlab 2021b (MathWorks) and the Circular Statistics toolbox 2012a (Berens, 2009).

#### Attentional manipulation

To ensure that attention was successfully manipulated (Carrasco, 2011; Dugué and VanRullen, 2014; Dugué et al., 2016, 2019; Senoussi et al., 2019), we assessed performance in the discrimination task as per d-prime (Hanning et al., 2019) (Eq1)

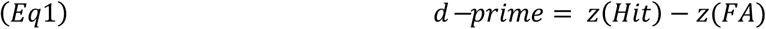

with *z*(*Hit*) and *z*(*FA*) as the *z* score for hit and false alarm rates. We also checked reaction times from Response 1 screen onset (**Figure 1**) to rule-out speed-accuracy trade-off (Wickelgren, 1977; Bogacz et al., 2010; Heitz, 2014; Huang et al., 2017).

**Figure 1.**
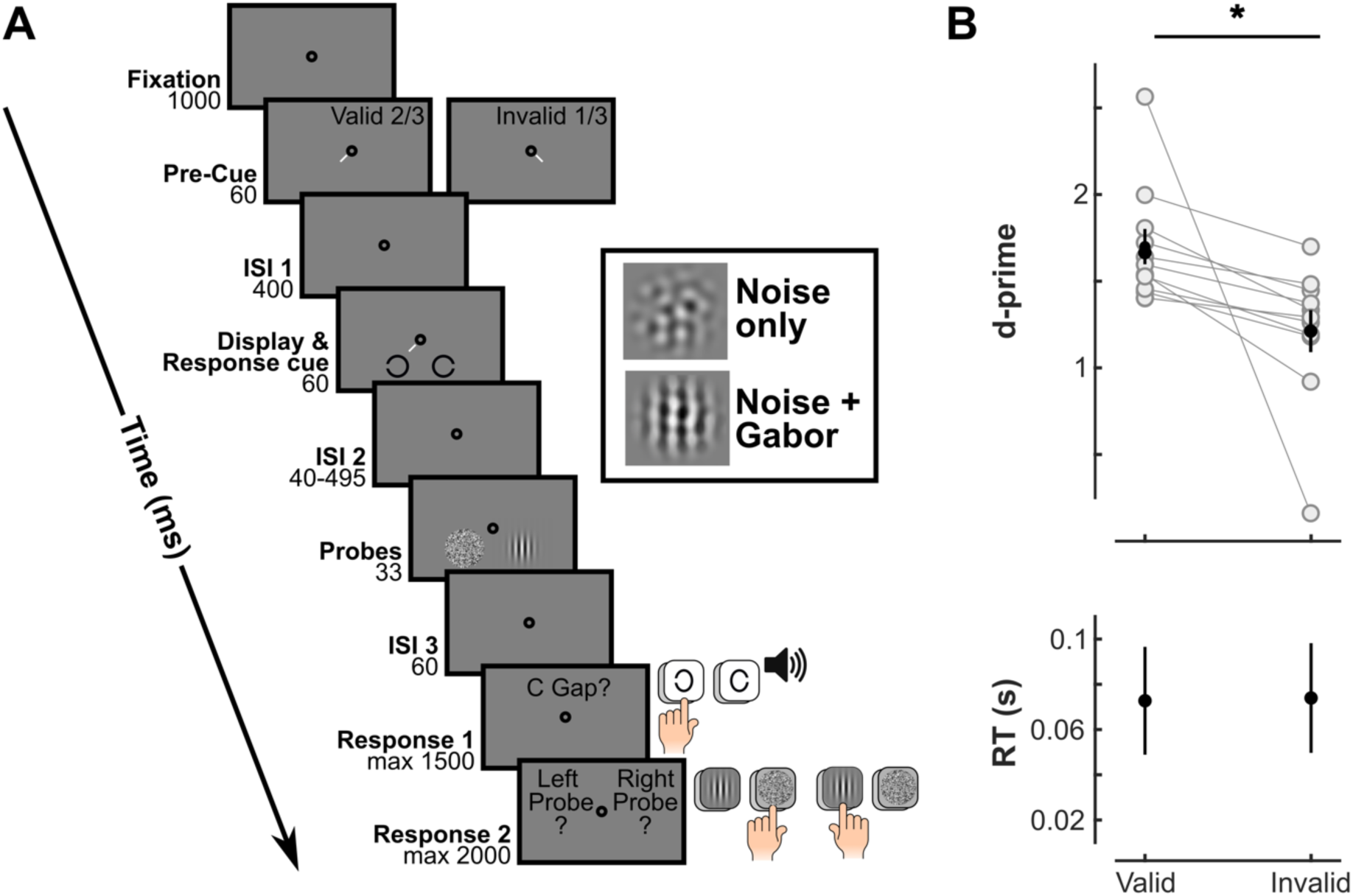
Attentional manipulation. **(A)** Trial Sequence. Trials began with a central fixation period of 1000ms. An endogenous pre-cue appeared for 60ms. After 400ms, Landolt-Cs were displayed for 60ms together with a response cue indicating which Landolt-C was the target (2/3 of valid trials, i.e., target at pre-cued location). Each Landolt-C could be oriented to the right or to the left randomly. After a variable delay from 40ms to 495ms (35ms step; only the fixation circle remained on the screen), two patches were presented for 33ms at the same location as the Landolt-C. Each patch could either show noise only (1/2 of trials) or noise + Gabor (1/2). After a third ISI of 60ms, participants were instructed to report: (1) the orientation of the target Landolt-C (right/left; discrimination task), and (2) the presence or absence of a Gabor in each of the two patches. **(B)** Behavioral results of the discrimination task. Individual (gray) and averaged (black) d-primes and median reaction times (RT) in the valid and invalid conditions. Error bars: standard error of the mean. *: significant difference between valid and invalid trials (p=0.003).

The validity effect (comparing performance between valid and invalid trials) was first assessed for each individual participant. D-prime was computed for each session and non-parametric statistics were performed: Wilcoxon Signed Rank test for paired-sample statistical comparison of valid and invalid d-prime. Out of the 15 participants, 4 participants did not show significant difference between valid and invalid trials, suggesting that the attentional manipulation was not successful. The following analyses were thus conducted on the remaining 11 participants. We report effect sizes (Eq2)

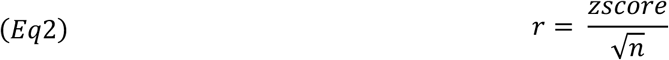

computed following Rosenthal recommendations (Rosenthal, 1994) with zscore from the Wilcoxon Signed Rank test and n the number of observations.

#### Behavioral rhythms

For the detection task, d-primes were examined for each cueing (valid and invalid) and ISI (delay between the Landolt-C and the probe patches) condition (Note that d-primes were normalized for visualization purposes only, **Figure 2**; all analyses were performed on non-normalized d-prime). Responses to both patches were combined in order to increase the number of trials for reverse correlation analysis and because we did not have a priori hypotheses regarding differences between the left and right locations. In total, there were 720 trials per ISI and participant. Fast Fourier Transform decompositions (FFT) were further performed on averaged d-primes to assess possible periodicity. Amplitude spectra for valid and invalid trials were plotted from 2.04Hz to 14.29Hz with steps of 2.04Hz (8 different frequencies). Phase angles were extracted from FFTs performed on each participants data for 12.2Hz in valid and 6.1Hz in invalid. We selected high- and low-performance trials by taking local maxima (above d-prime average) and minima (below d-prime average) for each of the d-prime time courses (ISI valid high: 145, 285, 390 and 460ms; low: 75, 180, 320, 425 and 495ms; ISI invalid high: 110, 320 and 425ms; low: 40, 145, 355 and 460ms). Two local maxima were removed, one in each condition, because they were below the averaged d-primes. All analyses described below were done separately for the high- and low-performance trials.

**Figure 2.**
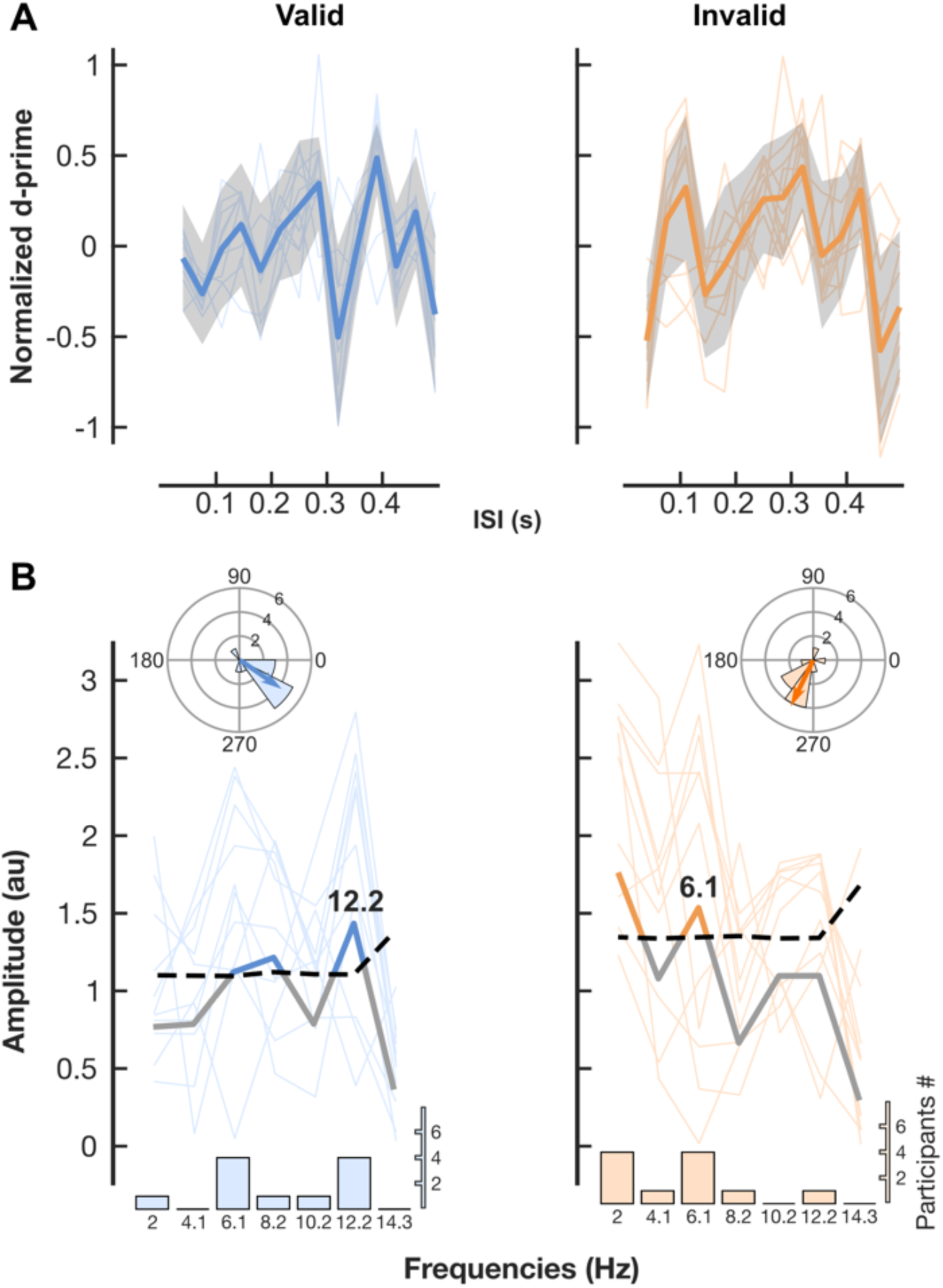
Behavioral rhythms. **(A)** Individual (light traces) and averaged (dark traces) d-prime time-courses (n=11) for valid (blue) and invalid (orange) conditions separately. Shaded area: 95% confidence interval. **(B)** Amplitude spectra obtained from FFT of individual (light traces) and averaged across participants (dark traces) d-primes time course. Dotted line: p=0.001 (permutation test for averaged d-prime time course). Polar histogram of individual phases (light bins) and weighted phase averages (arrows) at 12.2Hz in the valid condition and 6.1Hz in the invalid condition. Bar plots: number of participants at each frequency with the highest peak in individual amplitude spectra.

Note that valid and invalid conditions do not have the same number of trials. Also, the ISI selection detailed above shows differences between validity condition and high/low performance. To assess the potential impact of such differences in trial number, all previous analyses for both discrimination and detection tasks were also performed on subsampled trials (100 repetitions, random selection) thus equalizing trial numbers across all conditions. The results were the same as without subsampling (analysis not shown). We thus present the results from non-subsampled data.

All previous analyses were similarly performed on the decision criterion to assess whether the observed periodicity is exclusively on sensitivity or can also be observed on decision bias. Since in our task criterion is not independent of d-prime, we calculated criterion-prime (Eq3) was computed as follows

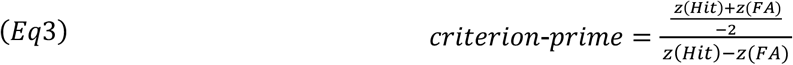

which consider d-primes variation to keep the decision boundaries stationary. No periodicity was observed in criterion-prime (analysis not shown).

#### Reverse Correlation analysis

To address behavioral feature tuning, we first transformed the noise from luminance intensity into the energy of different orientation components. We converted the noise image of each trial from pixel space (luminance intensity of each pixel) to a 1D space defined by the noise energy of components responding to different orientations (varied across all of orientation space, −90° to 90°, in steps of 7.5°, for 25 points on a linear scale). To compute the energy of each component, we took the noise image ***S*** of each trial, and convolved it with two grating filters (*g*_*θ*,*sin*_ in sine phase and *g*_*θ*,*cos*_ in cosine phase) with the corresponding orientation. The energy was computed as:

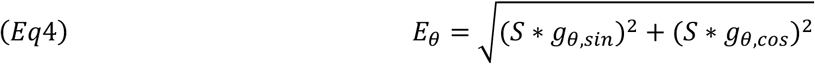

in which * represents the cross-correlation operator. We took the energy centered at the test stimulus (vertical orientation) for analysis. For each component with preferred orientation *θ*, we estimated the correlation between the energy of that component and behavioral responses using a GLM to predict the binomial dependent variable:

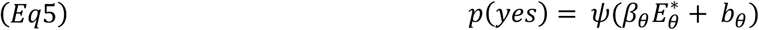

in which *p*(*yes*) was the percentage of yes responses in the detection task and *ψ* was a cumulative normal distribution. Two free parameters *β_θ_* and *b_θ_* were fitted. *β_θ_* represented the correlation between the energy and the behavioral response. A zero *β_θ_* indicated that the energy of that component did not influence participants’ responses. *b*_θ_ represented a baseline tendency of the participants to respond “present” (i.e., false alarm rate), which was not related to the stimuli. 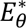 represented the centered and normalized energy of each component. Before applying the GLM, the energy of each component was first sorted into two groups based on whether the target signal was present or absent in each trial, and the mean of the energy was removed for each group. Subtracting the mean stimulus energy for target Gabor present and target Gabor absent trials independently serves two purposes: 1) it removes Gabor stimulus energy in present trials, so that the noise energy can be assessed independent of Gabor energy; 2) it centers stimulus energy for all trials to the mean, thus allowing straight-forward correlation of the responses (1 for present, −1 for absent) with stimulus energy – where positive values indicate greater-than-average stimulus and negative values indicate lower-than-average stimulus energy for each orientation channel. Reverse correlation aims to quantify which aspects of the stimulus during any given trial contribute to a higher or lower likelihood of reporting “present” irrespectively of the presence or the absence of a Gabor. Therefore, it is important to assess the difference of the noise on a given trial from the mean across all trials, and correlate that with behavior. If stimulus energy of a particular orientation is higher than average, and “present” responses are more likely, then the correlation is positive, and the opposite is true if high energy is associated with less likely “present” responses. To let the estimated *β*_!_ be comparable across components, we further normalized the energy across all the trials in each component to have a standard deviation of 1. The estimated sensitivity kernel was a 1D matrix *K* in which *K*(*θ*) = *β_θ_*. Before Gaussian fitting, positive and negative orientations’ energy were averaged.

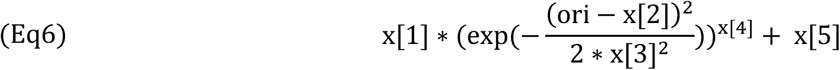

A Gaussian tuning curve was fit to the data (Eq6) for each condition, where ori represented all noise orientation channels (25 orientations). As we were agnostic as to the precise shape of the tuning function, in addition to the single Gaussian fit (Eq6) –a simple, standard model of feature tuning – a double Gaussian fit –a “Mexican hat” (Eq7) shape that sometimes arises due to attention (Müller et al., 2005)– was computed as well.

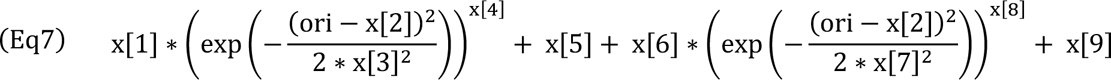

AIC corrected (AICc) for small data samples were performed using Python (3.8.1)(Anon, n.d.) independently for each condition to compare single and double Gaussian fits. AICc criterion was computed as shown in (Eq8) where k was the number of parameters, *L* the maximized value of the likelihood function and n the number of participants.

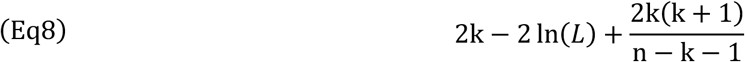

For each condition, the single Gaussian fit better explained the results (valid high-performance AICc difference (single fit – double fit) = −16.03; valid low-performance AICc difference = −37.34; invalid high-performance AICc difference = −37.34; invalid low performance AICc difference = −37.33). All further analyses were thus performed with the tuning curve from the single Gaussian fits.

We assessed the statistical differences between high- and low-performance trials’ tuning curves using linear mixed model with R (4.2.2), and specifically, differences in the standard deviation, amplitude and baseline of the tuning curves. Linear mixed models were fitted by maximizing the restricted log likelihood and using a normal distribution function. Performance was used as predictor with 2 levels (high and low) and participants as random effect. Non-parametric, paired-sample Wilcoxon Signed Rank tests were further performed to compare, for each orientation valid, high- and low-performance decision weights (FDR correction for multiple comparisons was applied).

Similar to the d-prime analyses, we assessed the potential impact of differences in trial number using a subsampling procedure (100 repetitions, random selection). The results were the same as without subsampling (analysis not shown). We thus present the results from non-subsampled data.

### Data and code availability

All data and analysis code are available on the OSF repository (https://doi.org/10.17605/OSF.IO/CXKBU).

## Results

### Successful attentional manipulation

We used a well-established psychophysics protocol (**Figure 1A**) tailored to identify rhythms in attentional tasks (for review (Kienitz et al., 2022)). Covert, voluntary attention was manipulated using a pre-cue (2/3 validity). Participants were instructed to first perform adiscrimination task – indicating the side of the gap, left or right, of a target Landolt-C – allowing us to confirm that attention was successfully manipulated and deployed to the pre-cued location. Only participants showing a significantly higher d-prime for valid (sustained mode of attention; target at the pre-cued location) than invalid (exploratory attention; target at the unpre-cued location) trials were included in the following analyses (see **Analyses and Statistics section**). The included participants are represented in **Figure 1B** (non-parametric statistics were used to describe the grand-averaged effect; Wilcoxon signed-rank test: z = 2.93, p-value = 0.003, r = 0.63; r>0.5 is considered a large effect size). Reaction times (RTs) were similar for valid and invalid trials (z = 1.42, p-value = 0.15, r = −0.30; note that RT values are short because they are measured from the onset of the Response 1 window), indicating no speed-accuracy trade-off (e.g., Dugué et al., 2018; Senoussi et al., 2019; Duyar et al., 2023).

### Attentional performance fluctuates periodically at 12.2Hz for sustained and 6.1Hz for exploratory attention

In the same trial, after the discrimination task and a variable delay, participants were presented with a second target, on which they were asked to perform a detection task reporting the presence or absence of a vertical Gabor embedded in band-passed noise (same protocol as in Li et al., 2016). The noise had random orientation content, drawn from a uniform distribution across all orientations −90 to +90 degrees (polar angle) and a spatial frequency of 3 cycles per degree (cpd). On half of the trials, a Gabor (3 cpd sinusoidal grating in a gaussian envelope subtending 2°; oriented vertically) was embedded in the noise patch. This second task was used to probe attention at each of the two stimulus locations and at various times during attentional orienting (valid trials; sustained attention) and reorienting (invalid; exploratory attention), assessing potential rhythms in attentional sampling.

We computed d-primes for the detection task independently for the valid and invalid conditions and for each ISI between the discrimination stimulus offset and detection probes onset. Across ISIs, d-prime exhibited multiple peaks and troughs, and individual data makes clear this was highly consistent across participants (**Figure 2A**; note that d-prime is normalized in the figure, but analyses were performed on non-normalized d-primes).

We then performed Fast-Fourier Transforms (FFT) of the averaged d-primes for the valid and invalid conditions separately. Peak frequencies were identified: 12.2Hz in valid and 6.1Hz in invalid (permutations tests: 100,000 surrogates, p = 0.001 after correction for multiple comparisons; note that we purposefully selected a conservative significance criterion to alleviate concerns regarding statistical power, Brookshire, 2022; **Figure 2B**). Secondary effects were observed in the valid condition at lower frequencies, indicating some variabilities across participants (see **Discussion** section). We concentrated the next analyses on the 12.2Hz effect, which was the most prominent effect and was consistent between the grand-averaged and individual FFT analyses. In the invalid condition, an additional significant peak frequency was observed at 2.04Hz reflecting the inverted U-shape trend (no detrending was performed here, see (Fiebelkorn et al., 2013; Michel et al., 2022); we did not analyze it further).

We further showed that the 12.2Hz (for the valid condition) and 6.1Hz (for the invalid) individual periodic modulations were in phase across participants (Rayleigh tests for circular data showed significant non-uniform distributions for both valid: Z = 6.14, p = 0.001; and invalid: Z = 4.54, p = 0.0075). Finally, in a supplementary analysis we examined whether there was periodicity in decision bias (criterion), as was found in sensitivity (d-prime). Because d-prime differed across ISIs, we needed to take that into account by controlling for d-prime in our measurement of decision bias. Specifically, we calculated criterion-prime (criterion ÷ d-prime (Macmillan and Creelman, 2005); measure of decision bias unaffected by fluctuations of d-prime), in both valid and invalid conditions. We found no periodicity in criterion-prime. Our results are thus not attributable to changes in decision bias (analysis not shown).

Together, our results replicate previously observed behavioral rhythms (Dugué and VanRullen, 2014; Dugué et al., 2016, 2019; Senoussi et al., 2019), and specifically show a highly consistent effect in sensitivity across participants (see also Li et al., 2016).

### Rhythmic sensory representations

For valid and invalid conditions separately, we split the ISI into low- and high-performance ISIs. ISIs corresponding to d-prime local maxima (and which were above the mean d-prime) were categorized as *high-performance* and ISIs corresponding to d-prime local minima (and which were below the mean) were categorized as *low-performance*. We then combined all high-performance trials (coming from 4 different ISIs in valid: 145, 285, 390 and 460ms; 3 ISIs in invalid: 110, 320 and 425ms) and low-performance trials (5 ISIs in valid: 75, 180, 320, 425 and 495ms; 4 ISIs in invalid: 40, 145, 355 and 460ms) for the next analysis. This trial separation was then used to assess modulations in sensory representations using reverse correlation analysis.

Fluctuations in sensitivity (d-prime) suggest a modification of sensory representations over time, such that high performance should be associated with more optimal sensory tuning compared to low performance. Here we assess which characteristics of sensory representations varied. Sensory tuning was modeled as a Gaussian distribution across orientations (**Figure 3**), thus tuning could vary via differences in: **(A)** tuning sharpness: for high-performance trials, high decision weights arise only for orientations more similar to the target orientation; **(B)** gain or amplitude: for high-performance trials, decision weights are multiplicatively higher across orientations; and **(C)** suppression or baseline shift: for high-performance trials, irrelevant orientations, i.e., those far from the target orientation, are suppressed more successfully.

**Figure 3.**
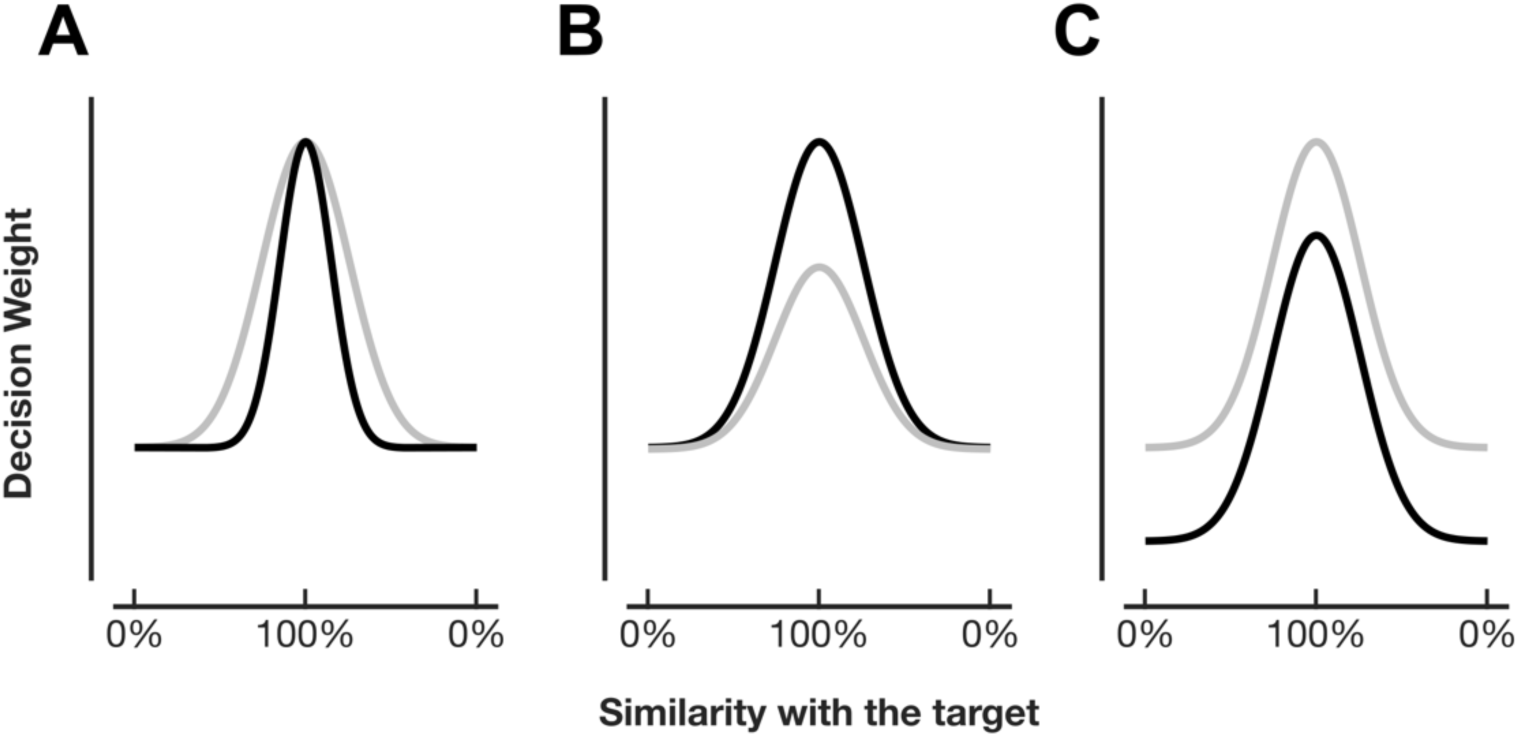
Sensory tuning hypotheses. Gaussian model of participant’s sensory tuning. Sensory representations are different during phases of high (black traces) versus phases of low (gray traces) performance. Modifications in the **(A)** tuning sharpness, **(B)** tuning amplitude (gain), **(C)** tuning baseline.

Reverse correlation of behavioral performance (see **Materials and Methods**) allowed testing of these three hypotheses, separately for each attention condition – sustained (valid) and exploratory (invalid). To infer the underlying sensory representation from the behavioral responses, we regressed the stimulus energy (i.e., visual resemblance of the noise to a range of orientation values) across trials –separately for high- and low-performance trials– with each participant’s responses (proportion of present vs. absent reports) using a binomial regression. This provided a sensitivity kernel for each orientation, i.e., β (or decision) weight, representing the level of correlation between the proportion of trials reporting “Target Present” and the energy of that particular orientation in the noise of the stimulus (**Figure 4**; Note that decision weights were averaged for positive and negative orientations). A high decision weight indicates a high correlation, suggesting that more energy for the orientation value in question within the noise positively influenced the likelihood of “Target Present” responses. The results show, first, that the highest decision weights were observed for orientation corresponding to the target grating orientation (0°; peak of the Gaussian). For each orientation, a paired-sample Wilcoxon Signed Rank test on the difference between high- and low-performance trials’ decision weights was then computed and FDR corrected for 13 comparisons (p-threshold FDR corrected = 0.0038; positive and negative orientation results were averaged). We observed significant differences for the values far from the target orientation mainly in the valid condition, and for near-target orientation values in the invalid condition.

**Figure 4.**
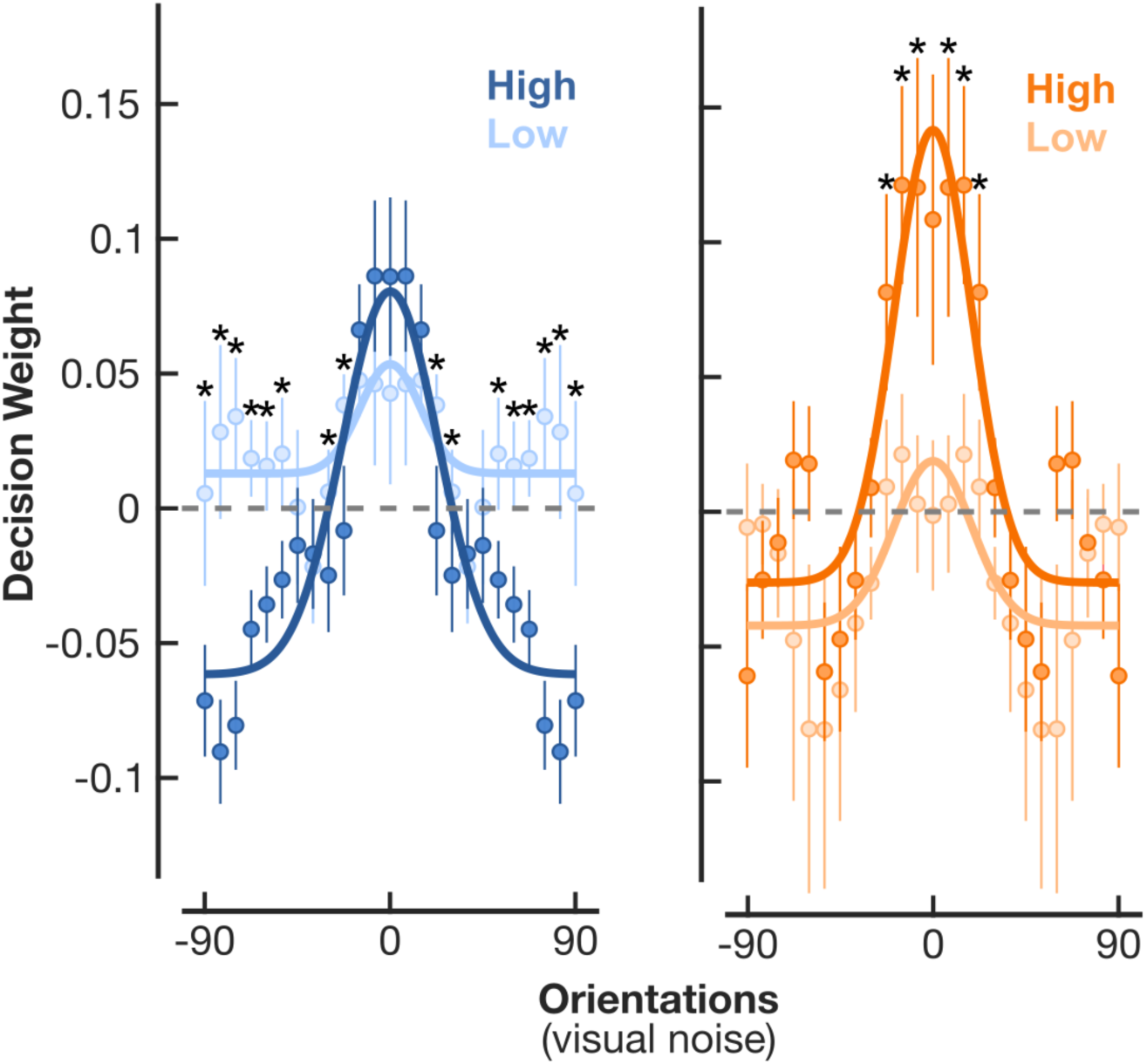
Rhythmic sensory representations: Decision weights from reverse correlation for valid and invalid trials. Dark traces: high-performance trials. Light traces: low-performance trials. Circles: average weight across participants of the correlation between energy at each orientation (averaged across positive and negative orientations) and participant’s response using a GLM. Error bars: 95% confidence interval. Solid lines: Gaussian fit. *: significant difference between high- and low-performance (p-threshold FDR corrected = 0.0038).

We then fit a single gaussian curve –the tuning curve– to the 25 decisions weights (1 per orientation; see **Materials and Methods**). To identify the changes in tuning that accompany periodicity in d-prime, we used linear mixed model to compare, across participants, each of the three following free parameters in the tuning fits: standard deviation (SD), amplitude and baseline, between high-performance and low-performance trials, separately for the valid and invalid conditions. There was no significant difference between high- and low-performance trials in both valid and invalid conditions for the SD (valid: t(10) =1.36, p = 0.205, estimate = 15.09 ± 11.13, standard error; invalid: t(10) = −0.197, p = 0.848, estimate = −2.73 ± 13.84).

Amplitude was significantly higher in high-compared to low-performance trials in both valid and invalid conditions (valid: t(10) = −6.57, p = 0.0001, estimate = −0.12 ± 0.02; invalid: t(10) = −3.94, p = 0.003, estimate = −0.11 ± 0.03). Baseline was significantly lower in high-compared to low-performance trials for the valid condition only (valid: t(10) = 7.62, p = 0, estimate = 0.08 ± 0.01; invalid: t(10) = −0.407, p = 0.692, estimate = −0.008 ± 0.02). These results indicate that, in the valid condition, when attention was sustained at one location, the performance fluctuation at alpha frequency was mediated by suppression of distracting features (baseline; hypothesis C) with a corresponding enhancement (gain) that kept sensitivity at target orientations constant across time (**Figure 3**); whereas, in the invalid condition, when attention had to explore the space, the fluctuation at theta frequency was mediated exclusively by a rhythmic change in enhancement (gain) of relevant features (**Figure 3**, hypothesis B).

## Discussion

The rhythmic sampling theory of attention has become an important topic of research yet remains widely debated due to methodological considerations and a lack of clear understanding regarding the sensory mechanism underlying performance fluctuations. We addressed this gap using a paradigm with a large amount of data collected for each participant and applied reverse correlation analysis that relates stimulus sensory information with behavioral performance. We assessed fluctuations in sensory representations for sustained (valid) and exploratory (invalid) attention. Attention was manipulated using a spatial cueing paradigm. Reaction times were short (measured after the long delay between target presentation and the response window), and below 600ms –typical time required for working memory processes to occur (Phillips, 1974).

Critically, performance sensitivity (d-prime; not criterion-prime) showed a main periodic fluctuation at the alpha frequency (12.2Hz) for sustained attention and theta (6.1Hz) for exploratory attention, replicating previous reports (Dugué et al., 2017, 2019; Senoussi et al., 2019; Michel et al., 2022; for review Dugué and VanRullen, 2017; Kienitz et al., 2022). Such effects were observed for each participant, with far more individual data than previous studies. This replication represents a strong confirmation of rhythmicity of voluntary attention, and the distinct frequency profiles of its sustained and exploratory modes. Interestingly, for valid trials, we found a large decrease in d-prime around 300ms. This effect has been observed in previous publications of behavioral rhythms (Dugué et al., 2015a; Dugué and VanRullen, 2017). We speculate that if behavioral sampling emerges from iterations between different cortical regions, 300ms may be the average time for the average number of iterations; further investigation is warranted. Note also that there were secondary effects at lower frequencies in the valid condition. They likely reflect harmonics, similar to the 12.2Hz non-significant peak in the invalid condition. We did not investigate inter-individual variability. In this study, we favored a high quantity of trials (10,080) per participant than a large sample based on recent recommendations (Smith and Little, 2018). It would be informative to replicate this study in independent samples.

Importantly, reverse correlation analysis was used to infer the neural representation underlying behavioral rhythms, i.e., to reveal the system’s sensitivity to and selectivity for stimulus feature (orientation), and approximate electrophysiological tuning properties. The results revealed differences in feature tuning across time between the two attention modes. Sustained attention displayed rhythmic suppression of distracting features –a change in baseline that decreased the influence of the most irrelevant orientation energy, and a corresponding change in gain that kept the influence of target orientation energy constant. Exploratory attention exclusively showed enhancement of relevant features.

Previous behavioral and electrophysiological evidence supports the rhythmic nature of perception and attention. Behavioral studies revealed rhythmic fluctuations of performance (reaction time, d-prime, accuracy) in various sensory modalities (vision (Landau and Fries, 2012; Fiebelkorn et al., 2013; Dugué et al., 2015a, 2017; Senoussi et al., 2019), audition (VanRullen et al., 2014; Ho et al., 2019), somatosensation (Baumgarten et al., 2015)) and types of behavioral response (finger, hand, arm, eye movements (Chota et al., 2018; Kienitz et al., 2018; Benedetto and Morrone, 2019)). It was suggested that when attention is sustained at one location, information is sampled at the alpha frequency, whereas, when exploring, attention samples information at the theta frequency (for review Dugué and VanRullen, 2017; Kienitz et al., 2022) – consistent with the results of the current study. This specific frequency effect was also reported in EEG studies (Dugué et al., 2015a; Merholz et al., 2022). As this line of research has emerged and matured, there has been a growing need to characterize the specific neural computations underlying periodic fluctuations in behavioral performance. Authors have speculated whether or not such variations were due to fluctuations in decision bias. It was observed that changes in EEG alpha activity could be explained by changes in decision criterion (Samaha et al., 2020). Here, using behavior alone, we demonstrated that fluctuations in sensitivity were not attributable to decision bias as there was no periodicity in the criterion-prime, for both the valid and invalid conditions. Tuning curves from reverse correlation analysis were used as proxy for neural computations (Ahumada, 2002; Eckstein et al., 2002; Paltoglou and Neri, 2012; Wyart et al., 2012; Li et al., 2016; Tu et al., 2023), taking into account the influence of specific participants’ responses.

Previous decades of attention research have concentrated on the effects of attention on perception and showed evidence for both signal enhancement and external noise reduction. Authors have demonstrated that attention can enhance signals through contrast gain (e.g., Lee et al., 1999; Cameron et al., 2002; Huang and Dobkins, 2005; Ling and Carrasco, 2006), response gain (e.g., Huang and Dobkins, 2005; Ling and Carrasco, 2006), orientation sensitivity gain (e.g., Lee et al., 1997, 1999), spatial resolution gain (e.g., Yeshurun and Carrasco, 1998, 1999; Carrasco et al., 2002; Golla et al., 2004), or information processing speed gain (e.g., Carrasco and McElree, 2001). Similarly, active suppression of irrelevant information occurs, also called external noise reduction. Interestingly, alpha-band brain oscillations seem to prominently contribute to this noise reduction, as evidenced in brain recordings (Foxe and Snyder, 2011; Händel et al., 2011; Bonnefond and Jensen, 2013; Wöstmann et al., 2019). Given the accumulated evidence for both attentional mechanisms (enhancement and suppression), they are almost certainly not mutually exclusive. Dosher and Lu (2000) indeed showed that either mechanism can arise in the same task under different levels of difficulty: stronger signal enhancement effect with low external noise; and stronger external noise reduction effect with higher external noise. They have also argued that which of the two mechanisms is engaged depends on the type of attention deployed (Lu and Dosher, 2000).

Here we found both signal enhancement and external noise reduction. When attention was directed to the target location, attention suppressed irrelevant features far from the target orientation (baseline shift), and then compensated with an increase in gain (amplitude). This resulted in equal decision weights at the target orientation for both high- and low-performance phases of d-prime sensitivity, but lower decision weights far from the target orientation at periods of high-compared to low-performance. In other words, when attention is oriented and sustained at a target location, alpha rhythms in performance are primarily due to fluctuations in the suppression of irrelevant features –advantageous suppression at peaks and poorer suppression at troughs– while a gain mechanism keeps the influence of relevant stimulus information constant over time. When attention is instead reoriented from an irrelevant location to a relevant one, theta rhythms in behavioral sensitivity are associated only with changes in the amplitude of the tuning function, most strongly influencing sensitivity around the target feature, such that high-performance phases have higher decision weights at orientations close to target feature compared to low-performance phases.

Notably, we found external noise reduction and signal enhancement in separate modes of voluntary spatial attention (sustained and exploratory), while difficulty remained constant. We speculate that the pattern of our findings is related to the distinct functional roles of the sustained and exploratory modes of attention. Because attention efficiently allocates resources based on task demands (Lennie, 2003; Carrasco, 2011; Merholz et al., 2022), finding distinct behavioral tradeoffs during different modes of attention suggests the two modes have distinct roles in aiding perception and performance. When attention shifts from one location to another, i.e., attention is exploring space, signal enhancement of the most relevant features may aid in locating the target, which is necessary before it can be examined and discriminated. When attention is sustained on a single location, i.e., the target location has already been identified, perception is enhanced via external noise reduction, as it is no longer necessary to find the target via signal enhancement.

The particular frequency of either sensory tuning modulation may be related to their corresponding attention modes’ respective functional roles (see reviews Dugué and VanRullen, 2017; Kienitz et al., 2022). Alternatively, either sensory mechanism may operate necessarily at a particular frequency due to metabolic or neural processing constraints. The link between attention mode, the respective frequencies of performance modulation, and the underlying sensory tuning mechanisms is a topic in need of further investigation.

Using a reverse correlation analysis of participants’ responses and trial-by-trial random noise information, we inferred the underlying neural mechanism. The reverse correlation tuning curves are comparable to those from neural recordings (Neri and Levi, 2006; Fernández et al., 2022). Critically, if the decision weights derived from reverse correlation are a true estimate of sensory weighting, then fluctuations of these weights can be logically related to fluctuations of neural representations. Such a statement is coherent with previous studies showing a link between rhythms of attentional performance and brain oscillations at the same frequencies (Dugué et al., 2015a; Fiebelkorn et al., 2018; Helfrich et al., 2018; Kienitz et al., 2018). Yet, due to the constraints imposed by such an intensive protocol, we do not have direct measurements of neural activity. This study will, however, provide valuable insights for further neurophysiological research.

## Acknowledgements

This project has received funding from the European Research Council (ERC) under the European Union’s Horizon 2020 research and innovation programme (grant agreement No 852139 - Laura Dugué). We also thank Laetitia Grabot, Kirsten Petras and Marisa Carrasco and her lab for their useful comments on the manuscript and advice on analyses.

